# Phytochemical studies and Antimicrobial Activity on the leaves of *Antigonon leptopus* and *Ecbolium viridae*

**DOI:** 10.1101/2024.04.16.589845

**Authors:** S Rudhra, A. Venkatesan

## Abstract

The aim of the present study is to investigate the in vitro Antibacterial activity of *Antigonon leptopus* and *Ecbolium viridae* methanol, ethanol, acetone leaf extracts for against selected bacterial strains respectively. Preliminary phytochemical screening of the leaf crude extract revealed the presence of alkaloids, flavonoids, phenols, taninns, carbohydrates, saponins. The evaluation of leaf powder was supported by physico-chemical analysis. The antibacterial activities of leaf of *E*.*viride* and *Antigonon leptopus* in successive different solvent were tested against gram +ve (*Staphylococcus aureus, Streptococcus pyogenes*) and gram −ve (*Escherichia coli, Pseudomonas aeruginosa*) Antibacterial activity of those crude extracts are determined by using agar well diffusion method and the zone of inhibition of microorganisms was measured in mm. Amphicillin is used as a positive control respectively.

## 1. Introduction

Medicinal plants are still invaluable source of safe, less toxic, lower price, available and reliable natural resources of drugs all over the world. People in Sudan and in other developing countries have relied on traditional herbal preparations to treat themselves. Therefore, it is useful to investigate the potential of local plants against these disabling diseases (Amaral *et al*., 2006; Koko *et al*.,2008).It is commonly grown in gardens and often run wild. It is a climbing vine; stems slender. Leaves are alternate, cordate-ovate or triangular, entire, acute to acuminate. Flowers are bright pink, in panicled racemes that terminate in tendril. Fruits of 1-seeded, hard nut let, 3-gonous, biconvex, compressed (Madhava Chetty *et al*.,2008). Traditionally, *A. leptopus* have been used to treat diabetes, asthma, liver and spleen disorders, cough and throat constriction (Cheryl A Lans, 2006; Idu and Onyibe, 2007; Mitchell and Ahmad, 2006). The present study was conducted to investigate the *in-vitro* antioxidant (DPPH assay), phytochemical screening and cytotoxicity (MTT) of ethanol extract of *A. leptopus* (leaves) in Sudan. Plants can synthesize a large variety of chemical substances that are of physiological importance Many plants contain antioxidant compounds and these compounds protect cells against the free radicals. [Padmaja et al.,2011] During cell metabolism free radicals are produced in the cells, majorly through mitochondrial oxidative phosphorylation. Free radicals like reactive nitrogen species (RNS) and reactive oxygen species (ROS) (i.e., oxygen species such as singlet oxygen, superoxide, peroxyl radicals, hydroxyl radicals and peroxynitrite) are being produced. [Nadia *et al*.,2015] RNS are generated from oxygen in the presence of nitric oxide synthetase. ROS and RNS can be harmful as they damage cellular lipids, sugars, proteins, and nucleic acids, thus inhibiting the normal function. Oxygen free radicals have been shown to be responsible for many pathological conditions in human beings such as atherosclerosis, ischemic heart disease, aging process, inflammation, and diabetes. Thus, there is a continuous requirement of antioxidants for free radical inactivation.[Mathew *et al*.,2013] The extracted and purified methanolic extract of *Antigonon leptopus* yields n-hentriacontane, ferulic acid, 4-hydroxycinnamic acid, quercetin-3-rhamnoside, and kaempherol-3-glucoside along with sitosterol, b-sitosterol-glucoside and d-mannitol and hence shows as a antibacterial, anti-inlamatory, anti-oxidant activity, etc.[Mulabagal *et al*., *2008*] Also it has been used to treat diabetes, asthma, liver and spleen disorders, cough and throat constriction, flu-related pains, hypertension, antithrombin agent and used to reduced menstrual pains.[Cheryl Lans,2006; Chistokhodova *et al*.,*2002*] Literature survey revealed that the extract of plant was evaluated for structura characterization, antioxidant and anticancer properties of gold nanoparticles of extract,. [Balasubramani *et al*., *2015*] for new steroidal saponin,[Apaya MK, Chichioco-Hernandez CL.,2014] as well as for extermination of fish bacteria pathogens. [Balasubramani *et al*., *2015*]

## 2. MATERIALS AND METHODS

### 2.1. Collection of plant leaves

Leaves of *A*.*leptopus* and *E*.*viridae* are collected in annamalai nagar, Annamalai University, Chidambaram, Cuddalore District. Plants were appropriately rinsed with distilled water to purge dust, dirt and other possible parasites and then were shade dried at 25-30°C. The dried parts root, stem, and leaves were pulverized incoherently and then stored in clean, dried plastic bags for extraction.

### 2.2. Extraction of plant materials

For the preliminary phytochemical analysis, extract was prepared by weighing 50gm of the dried powdered leaf was subjected to hot successive continuous extraction with 250ml,methanol and aqueous extractions. The extraction process was carried out for 72hrs and then the extract was collected. It was evaporated in a hot plate and residue was collected, kept in desiccators. All the extracts obtained by successive extraction method are subjected to qualitative phytochemical analysis.

### 2.3. Qualitative phytochemical screening

Phytochemical screening of the plant samples was carried out using described protocols of Harbone, Sofowora, Trease and Evans with minor modifications. A stock solution of each extract, with a concentration of 10 mg extract/ml distilled water, was prepared and used for the

#### Phytochemical screening

The different chemical tests were performed for establishing profile of the leaf extract for its chemical composition. The chemical tests for various phyto-constituents in the crude extract was carried out as described below:

#### 2.3.1. Test for the presence of alkaloids

Approximately 3 ml of extracts were added to 3 ml of 1% HCl and heated for 20 min. The mixtures were then cooled and used to perform the following tests:

##### Mayer’s test

To the 1ml of plant extract in a test tube, 1 ml of Mayer’s reagent was added drop by drop. The formation of a greenish color or cream precipitate indicated the presence of alkaloids.

##### Dragendoff’s test

To the 1ml of plant extract in a test tube, 1 ml of Dragendoff’s reagent was added drop by drop. The formation of a reddish-brown precipitate indicated the presence of alkaloids.

##### Wagner’s test

To the 1ml of plant extract in a test tube, 1 ml of Wagner’s reagent was added drop by drop. The formation of a reddish-brown precipitate indicated the presence of alkaloids.

#### 2.3.2. Test for the presence of saponins

About 3 ml of plant extract was added to 3 ml of distilled water and shaken vigorously. The formation of a stable persistent froth was taken as a positive test for saponins.

#### 2.3.3. Test for the presence of flavonoids

About 2 ml of the plant extract was dissolved in 2 ml of methanol and heated. A chip of magnesium metal was added to the mixture followed by the addition of a few drops of concentrated HCl. The occurrence of a red or orange coloration was indicative of the flavonoids.

#### 2.3.4. Test for the presence of tannins

About 2 ml of plant extract was diluted in 5ml distilled water in a test tube, and then added a few drops of 1% lead acetate solution. A red precipitate shows the presence of tannins.

#### 2.3.5. Test for the presence phenols

Two ml of 5% solution of FeCl_3_ was added to 1 ml crude extract. A black or blue-green color indicated the presence of tannins and phenols.

#### 2.3.6. Test for the presence of terpenoids

Approximately 2 ml of chloroform and 3 ml of H2O2 were added to 5 ml of plant extracts. A reddish-brown coloration was taken as positive test for terpenoids.

### 2.4. Antibacterial activity

#### 2.4.1. Kirby-Bauer’s Disc diffusion method

Mueller Hinton agar plates were spread with 100 μl of actively growing broth cultures of the respective bacteria and are allowed to dry for 10 minutes. The sterile readymade discs loaded with each extract individually (100 μl/disc, 75 μl/disc and 25 μl /disc) were imposed on the inoculated plates. The plates were then incubated at 37°C for 36 hours. The development of the inhibition zone around the extract loaded disc was recorded. Sterile disc with respective solvent of 25 ml was used as negative control and Amphicillin at 10mg/disc was used as positive control.

#### 2.4.2. Bacterial cultures

The test organisms are gram positive (*Staphylococcus aureus, Streptococcus pyogenes* and *Leuconostoc lactis*) and gram negative (*Escherichia coli, Pseudomonas aeruginosa* and *Salmonella typhi*) from Department of botany, Annamalai University.

## 3. Result and Discussion

### 3.1. Phytochemical Analysis results

The present study revealed that the various alcoholic and aqueous extract of leaf of *Antigonon leptopus* contained alkaloids, flavonoids, terpenoids, triterpenoids, phenols, saponin and steroids. Except saponin other secoundary metoblites were detected in acetone extract of leaf parts. Next to acetone extract, ethanol extract of leaf parts showed the presence of rich variety of secondary metabolites. Methanol extract showed the less variety of these secondary metabolites. Compared to other solvent extract ethanol leaf extract had higher number of secondary metabolites with higher degree of precipitation. Mid polarity solvent of ethanol has ability to extract secondary metabolites from *Antigonon leptopus*. It responds to ethanol very much than other two solvents of acetone and methanol. Saponins are absent in two solvents on leaf parts of *A*.*leptopus. Ecbolium viridae* contained alkaloids, flavonoids, terpenoids, triterpenoids, phenols, saponin and steroids. Except saponin other secoundary metoblites were detected in acetone extract of leaf parts. Next to acetone extract, ethanol extract of leaf parts showed the presence of rich variety of secondary metabolites. Methanol extract showed the less variety of these secondary metabolites. Compared to other solvent extract ethanol leaf extract had higher number of secondary metabolites with higher degree of precipitation. Mid polarity solvent of ethanol has ability to extract secondary metabolites from *E*.*viridae*. It responds to ethanol very much than other two solvents of acetone and methanol. Saponins are absent in all three solvents on leaf parts of *E*.*viridae*.

### 3.2. Quantitative determination of the chemical constituency

#### 3.2.1. Extraction yield

Table 2.shows the percentage of yield of crude successive extract (methanol, acetone and Ethanol) of leaf parts of *A*.*leptopus and E*.*viride*. Ethanolic extracts of leaf exhibited higher yield of 5.07% and 5.87% followed by Acetone leaf extract showed the lowest yield of 3.65% and 4.09%. The methanol extracts showed the moderate amount of yield of 4.60% and 4.99%.

**Table 1.**
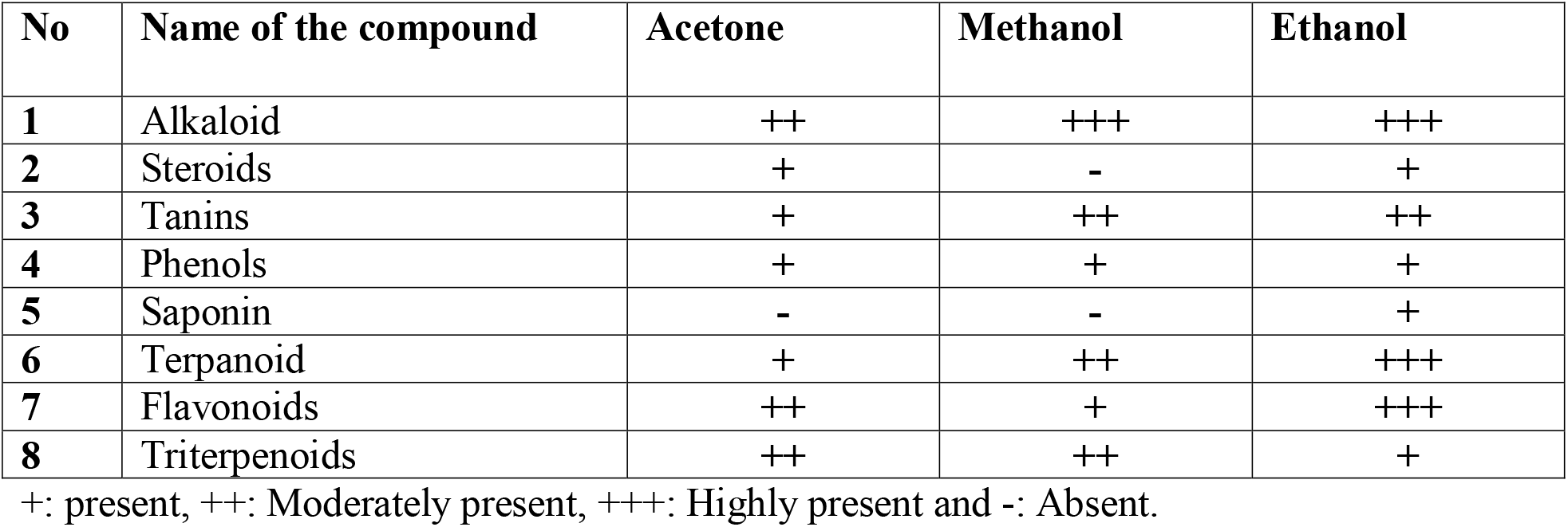
Preliminary qualitative phytochemical analysis of various extracts of leaf of *Antigonon leptopus*.

| No | Name of the compound | Acetone | Methanol | Ethanol |
| --- | --- | --- | --- | --- |
| 1 | Alkaloid | ++ | +++ | +++ |
| 2 | Steroids | + | - | + |
| 3 | Tanins | + | ++ | ++ |
| 4 | Phenols | + | + | + |
| 5 | Saponin | - | - | + |
| 6 | Terpanoid | + | ++ | +++ |
| 7 | Flavonoids | ++ | + | +++ |
| 8 | Triterpenoids | ++ | ++ | + |

+: present, ++: Moderately present, +++: Highly present and -: Absent.

**Table 2.** Preliminary qualitative phytochemical analysis of various extracts of leaf of *Ecbolium viridae*.

| No | Name of the compound | Acetone | Methanol | Ethanol |
| --- | --- | --- | --- | --- |
| 1 | Alkaloid | ++ | ++ | +++ |
| 2 | Steroids | + | - | ++ |
| 3 | Tanins | + | + | + |
| 4 | Phenols | + | + | ++ |
| 5 | Saponin | - | - | - |
| 6 | Terpanoid | + | + | + |
| 7 | Flavonoids | + | ++ | +++ |
| 8 | Triterpenoids | + | - | + |

+: present, ++: Moderately present, +++: Highly present and -: Absent.

#### 3.2.2. Qualitative test results

#### 3.2.3. Total phenol content

Total phenol content of various extracts of leaf of *A*.*leptopus* was varying widely between 0.094 to 2.565 mg GAE/100g extract (Table 3). Ethanolic extract of leaf were demonstrating higher total phenol content of 2.565 mg GAE/100g(Table 4) extracts than that of the other solvent extracts. Same goes for *E*.*viridae* also varying range between 1.054 to 3.089 mg GAE/100g extract. Ethanolic leaf extract is higher than other two solvents respectively.

**Table 3.**
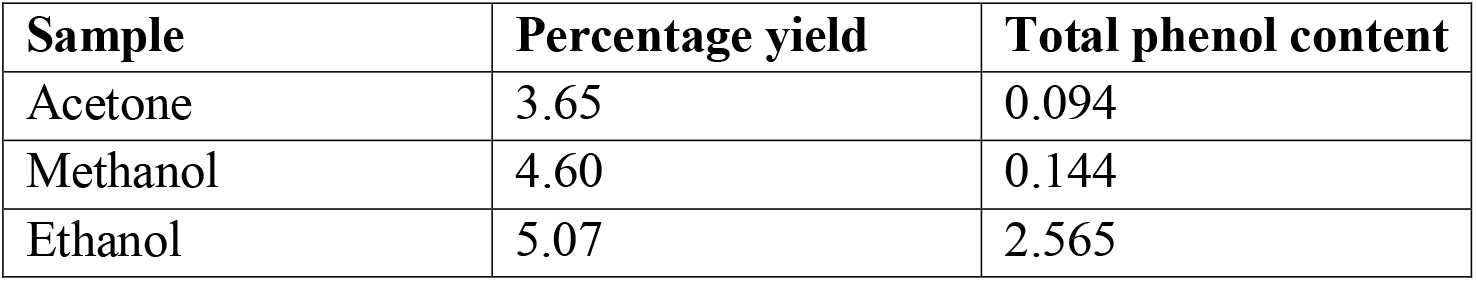
Percentage yield, total phenol contents of various alcoholic and aqueous extracts of leaf part of *Antigonon leptopus*.

**Table 4.** Percentage yield, total phenol contents of various alcoholic and aqueous extracts of leaf part of *Ecbolium viridae*.

| Sample | Percentage yield | Total phenol content |
| --- | --- | --- |
| Acetone | 4.09 | 1.054 |
| Methanol | 4.99 | 2.890 |
| Ethanol | 5.87 | 3.089 |

### 3.3. Antibacterial activity

The results showed that antibacterial activities of leaf extract of *A*.*leptopus* and *E*.*viridae* is presented in table 5. This antibacterial activity was quantitatively determined by the presence or absence of inhibition zone around the discs containing extract. The ethanol extract shows the highest antibacterial activity at 200(μg\mL) with inhibition zone of 20.4±0.13mm in *A*.*leptopus* and 20.3±0.27mm in *E*.*viriade* against *Staphylococcus aureus* followed by inhibition zone of 20.2±0.11mm in *A*.*leptopus and* 19.3±0.26 mm in *E*.*viridae* against *E. coli* and inhibition zone of 18.9±0.19 mm in *A*.*leptopus and* 16.56±0.3mm in *E*.*viridae* against *K. pneumonia*. The methanol extract exhibited moderate antibacterial activity at 200(μg\mL) with the inhibition zone of 16.1±0.08 mm in *A*.*leptopus and*13.2±0.12mm in *E*.*viridae* against *E. coli* and inhibition zone of 18.1±0.14 mm in *A*.*leptopus and* 19.3±0.26mm in *E*.*viridae* against *S. aureus* and inhibition zone of 13.2±0.12 mm in *A*.*leptopus and* 16.3±0.05mm in *E*.*viridae* against *K. pneumonia*. The lowest antibacterial activity was recorded in acetone extract with the inhibition zone of 12.5±0.12 mm in *A*.*leptopus* 11.8±0.53mm in *E*.*viridae* at 200 g/mL against *K. pneumonia* followed by inhibition zone of 12.4±0.11 mm in *A*.*leptopus and* 15.2±0.72 mm in *E*.*viridae* against *E. coli* and inhibition zone of 12.3±0.18 mm in *A*.*leptopus and* 18.5±0.16 mm in *E*.*viridae* against *S. aureus*. The most resistance strains among 3 tested bacterial species is *S. aureus*. In fact, the crude extract had not significant antibacterial effect against *K. pneumoniae*.

**Table 5.** Antibacterial activity of different crude extracts of *A*.*leptopus* and *E*.*viridae* leaf against pathogenic microorganisms using agar well diffusion method. Values are shown mean ± SD.

| Sl. No. | Microorganisms/<br>Solvents Extracts | Mean zone of inhibition* (mm) |  |  |  |  |  |
| --- | --- | --- | --- | --- | --- | --- | --- |
| | | Concentration of leaf extract (μg \ mL) | | | | | |
|  |  | 25 | 75 | 100 | 150 | 200 | Control |
| 1 | *A.leptopus*<br>*Escherichia coli* |  |  |  |  |  |  |
| | Acetone | 7.1±0.04 | 8.4±0.28 | 11.2±0.19 | 11.6±0.15 | 12.4±0.11 | 27.1±0.25 |
| | Methanol | 10.7±0.38 | 12.4±0.16 | 15.5±0.22 | 16.5±0.15 | 16.1±0.08 | 27.5±0.15 |
| | Ethanol | 12.2±0.16 | 14.1±0.39 | 17.4±0.13 | 18.0±0.07 | 20.2±0.11 | 28.7±0.15 |
| 2 | *Staphylococcus aureus* |  |  |  |  |  |  |
| | Acetone | 8.5±0.25 | 9.3±0.38 | 11.0±0.27 | 11.4±0.25 | 12.3±0.18 | 19.2±0.58 |
| | Methanol | 10.1±0.18 | 11.6±0.17 | 13.7±0.18 | 14.7±0.18 | 18.1±0.14 | 18.7±0.12 |
| | Ethanol | 11.4±0.20 | 13.5±0.15 | 16.8±0.21 | 18.1±0.14 | 20.4±0.13 | 19.9±0.35 |
| 3 | *Klebsiella pneumonia* |  |  |  |  |  |  |
| | Acetone | 9.7±0.30 | 10.3±0.22 | 11.2±0.11 | 11.5±0.25 | 12.5±0.12 | 19.1±0.25 |
| | Methanol | 9.8±0.13 | 10.6±0.16 | 11.8±0.35 | 12.2±0.22 | 13.2±0.12 | 18.2±0.15 |
| | Ethanol | 11.3±0.22 | 13.3±0.15 | 15.7±0.13 | 17.3±0.15 | 18.9±0.19 | 19.2±0.28 |
| 1 | *E.viridae*<br>*Escherichia coli* |  |  |  |  |  |  |
| | Acetone | 9.6±0.13 | 10.8±0.28 | 11.2±0.11 | 11.6±0.29 | 15.2±0.72 | 18.5±0.14 |
| | Methanol | 10.4±0.12 | 10.6±0.15 | 11.8±0.53 | 12.5±0.42 | 13.2±0.12 | 20.7±0.18 |
| | Ethanol | 11.3±0.22 | 13.3±0.15 | 15.7±0.13 | 17.3±0.15 | 19.3±0.26 | 20.6±0.19 |
| 2 | *Staphylococcus aureus* |  |  |  |  |  |  |
| | Acetone | 8.4±0.14 | 8.8±0.17 | 9.2±0.35 | 9.4±0.30 | 18.5±0.16 | 19.5±0.27 |
| | Methanol | 10.4±0.12 | 10.6±0.15 | 11.8±0.53 | 12.8±0.45 | 19.3±0.26 | 20.1±0.12 |
|  | Ethanol | 11.3±0.22 | 13.3±0.15 | 15.7±0.13 | 19.1±0.15 | 20.3±0.27 | 21.7±0.22 |
| **3** | ***Klebsiella pneumonia*** |  |  |  |  |  |  |
|  | Acetone | 7.76±0.11 | 8.46±0.11 | 10.42±0.23 | 10.67±0.13 | 11.8±0.53 | 19.5±0.35 |
|  | Methanol | 9.6±0.07 | 9.49±0.19 | 11.45±0.17 | 11.81±0.03 | 16.3±0.05 | 20.5±0.18 |
|  | Ethanol | 10.67±0.13 | 10.27±0.19 | 12.34±0.18 | 14.67±0.34 | 16.56±0.3 | 21.2±0.35 |

## 4. Discussion

Antibiotics have shown incalculable mental and material value in saving lives. However, along with the antibiotic era, a new threat called antimicrobial resistance emerged, which currently limits the successful completion of the centenary of the antibiotic era [Rahman, M.; Sarker, S.D, 2020; De Oliveira *et al*., *2020*]. The current role of scientists around the world is to meet the challenge of discovering new sources of effective antimicrobial drugs or to design and synthesize them. Medicinal plants have been the most valuable source of molecules with therapeutic potential throughout the history of mankind. Folk medicine of each civilization is based on natural products and nowadays, medicinal plants still represent an important pool for the identification of novel drug leads [Atanasov, 2015]. The trend of natural products for therapeutic use may be especially beneficial in the treatment of skin and wound infections, due to a good accessibility of these infected lesions for topical drug application. Skin with its protective barrier role and sensory, secretory, and thermoregulatory functions is the largest organ of the human body [Buchvald, 2019]. The increase of antimicrobial resistance against commercial drugs has led to the study of plant products for searching new antimicrobials [Clardy *et al*., *2006*]. Higher plants mainly produce antimicrobial compounds for their defense mechanism against infections. Their secondary metabolites or phytochemicals represent a large reservoir of structural moieties which work together exhibiting a wide range of biological activities. Phytochemicals act as antimicrobial agents by inhibiting peptidoglycan synthesis, damaging microbial membrane structures and modifying bacterial membrane surface hydrophobicity [Rasooli *et al*., *2008*]. Plants have many phytochemicals with various bioactivities, including antioxidant, anti-inflammatory, antimicrobial and antidiabetic activities. It can be observed from above results that inhibition was high against Gram positive bacteria than Gram negative bacteria due to the presence of an effective permeability barrier in Gram negative bacteria, comprising of an outer membrane with high content of cell wall lipopolysaccharides, which restricted the penetration of phytochemical amphipathic compounds and multi drug resistance pumps that extrude toxins across the barrier [Jaya raju N, Gangarao battu, 2009]. The reason for high MIC values exhibited by the crude extract might be because the active principle compound was present in very low concentration and also the active compound was present along with other secondary metabolites which might restrict the inhibitory activity against microbial strains in the crude extract.
The results of antimicrobial activity on *A*.*leptopus* and *E*.*viridae* were in acceptance with these reports. Correlation between the phytoconstituents and the bioactivity of plant has been established long back and identification of phytochemicals is desirable so as to isolate compounds with specific activities from the plant extract which can be used to treat various health ailments and chronic diseases as well [Pandey *et al*., *2013*]. Owing to the significance in the above context, such preliminary phytochemical screening of plants is the need of the hour in order to discover and develop novel therapeutic agents with improved efficacy. Numerous research groups have also reported such studies throughout the world [. Kharat *et al*., *2013*: Adesokan *et al*.,*2007*].

## 5. Conclusion

In recent years ethno medicinal studies received much attention on natural resources to light the numerous medicines, especially of plant origin which needs evaluation on modern scientific lines such as phytochemical analysis, pharmacological and clinical trials. Based on these results, it can be concluded that, the different solvents of leaves extracts of *Antigonon leptopus* and *Ecbolium viride* exhibited a broad range of antimicrobial activity to varying degrees. Particularly, methanol and acetone crude extract showed significant antimicrobial activity and could be used as an antimicrobial agent in new drugs for therapy. So, further evaluation and analysis of phytochemical compounds are needed in order to utilize them as novel antimicrobial agents. Hence, the present study justifies the claimed uses of *A*.*leptopus* and *Ecbolium viride* in the traditional system of medicine to treat various infectious diseases caused by microbes.

